# *Rēs ipSAE loquuntur*: What’s wrong with AlphaFold’s *ipTM* score and how to fix it

**DOI:** 10.1101/2025.02.10.637595

**Authors:** Roland L. Dunbrack

## Abstract

AlphaFold’s *ipTM* metric is used to predict the accuracy of structural predictions of protein-protein interactions (PPIs) and the probability that two proteins interact. Many AF2/AF3 users have experienced the phenomenon that if they trim full-length sequence constructs (e.g. from UniProt) to the interacting domains (or domain+peptide), their *ipTM* scores change, even though the structure prediction of the interaction is unchanged. The reason this happens is due to the mathematical formulation of *ipTM* in AF2/AF3, which scores the interactions of whole chains. When one chain is a single ordered domain but the other contains lots of disorder or irrelevant domains, *ipTM* is higher in the presence of those residues than it is when they are trimmed. If both chains in a PPI complex contain large amounts of disorder or accessory domains that do not form the primary domain-domain or domain/peptide interaction, the *ipTM* score can be significantly lower in the presence of those regions than when they are trimmed. The score then does not accurately represent the accuracy of the structure prediction nor whether the two proteins actually interact. We have solved this problem by: 1) including only residue pairs in the *ipTM* metric that have good predicted aligned error (*PAE*) scores; 2) by adjusting the *d*_0_ parameter (a function of the length of the query sequences) in the TM score equation to include only the number of residues with good interchain *PAE*s to the aligned residue; and 3) using the *PAE* value itself and not the probability distributions over the aligned error to calculate the pairwise residue-residue *pTM* values that go into the *ipTM* calculation. The first two are crucial in calculating high *ipTM*s for domain-domain and domain-peptide interactions even in the presence of many hundreds of residues in disordered regions and/or accessory domains. The third allows us to require only the common output json files of AF2 and AF3 (including the server output) without having to change the AlphaFold code and without affecting the accuracy. It also works on npz file output from Boltz. We show in a benchmark that the new score, called *ipSAE* (interaction prediction Score from Aligned Errors), is able to separate true from false complexes more efficiently than AlphaFold2’s *ipTM* score. The resulting program is freely available at https://github.com/dunbracklab/IPSAE.

## Introduction

The AlphaFold programs (1-3) have had a profound impact on the structure prediction of proteins and protein complexes. AlphaFold-Multimer (v2.3) has enjoyed the widest use in predicting the structures of protein-protein interactions (PPIs), which are critical to essentially all biological processes. Since AlphaFold-Multimer code has been available for download since late 2021 (and v2.3 since December 2022), these programs have been extensively benchmarked for their ability to predict the structures of protein complexes accurately and their ability to predict whether two proteins interact. These benchmarks have utilized the scoring output from the AlphaFold programs, including residue-specific predicted local distance difference tests (*pLDDT*s), predicted aligned errors (*PAE*s) for residue pairs, and predicted template-modeling scores (*pTM*) and interface predicted template modeling scores (*ipTM*s) for the whole modeled system.

Typically, benchmarks have been constructed from Protein Data Bank (PDB) structures, and use the sequences provided for each PDB entry (e.g., the CASP competitions (4, 5) and others (6, 7)). That is, they do not use the full UniProt sequences, which may contain disordered sequences and domains that do not form part of the interaction. PDB constructs are mostly fully ordered, save for some loops or short N and C terminal tails. In these cases, the *ipTM* score generally works well in assessing the accuracy of the structure prediction (8). However, in real-world situations where the interacting regions may not be known, structure predictions usually start with full-length protein sequences from UniProt. Then after observing which domains interact in the model with good *PAE* scores, it can be productive to input shorter sequence constructs to AlphaFold.

Many studies have noted that different sequence constructs produce different *ipTM* scores, even though the predicted interface contacts are unchanged (9-13). For example, Danneskiold-Samsøe et al. compared AlphaFold-Multimer v2.2 models produced from either full-length sequences of single-pass transmembrane receptors and their full-length unprocessed ligands, or various truncations of the proteins (e.g., the extracellular domains only and the proteolytically processed secreted ligand proteins) (14). *ipTM* scores were higher and more predictive for the shorter constructs comprising only the interacting domains. In a comprehensive study, Lee et al. found shorter fragments of peptides binding to protein domains often scored better than longer fragments or full-length proteins (15). Bret et al. developed a scanning approach to search through disordered sequence regions for protein domain binders, because the *ipTM* score was not successful on full-length sequences (9). Some reports have shown that *ipTM*s, which are calculated over whole chains, are less predictive than other measures. These measures include *pLDDT* values of interface residues (as in the pDockQ score) (16, 17), interface *PAE* values (*iPAE*’s) calculated over only interchain residue pairs within various cutoff distances (18, 19), or combinations of AlphaFold metrics and energy functions to evaluate interfaces (7).

In this paper, we investigate the origin of the behavior of AlphaFold’s *pTM* and *ipTM* scores based on their mathematical descriptions in the AlphaFold papers. We then use this analysis to identify alternative formulations that are not sensitive to disordered regions or non-interacting accessory domains in either or both chains of pairwise interactions in AlphaFold models. We show that using only interchain residue pairs with good *PAE* scores in the evaluation of *ipTM* and evaluating the TM formula’s *d*_0_ parameter (which is based on sequence length) according to the number of such pairs, we can produce good *ipTM*-like values for true interactions even in the presence of large amounts of disorder and accessory domains. The resulting code, which is freely available, works only on the *PAE* matrix provided in the default output of both AlphaFold2 and AlphaFold3. We have named the metric *ipSAE* for “interaction prediction score from aligned errors.” The word is a play on the Latin phrase “*Rēs* ***ipsae*** *loquuntur*,” meaning “The things speak for **themselves**,” referring to the AlphaFold output scores.

### Derivation of the *ipTM* and *ipSAE* scores

The TM score was developed by Zhang and Skolnick to assess the accuracy of predicted models of protein structures compared to experimental structures of the same proteins (20). It is defined as:

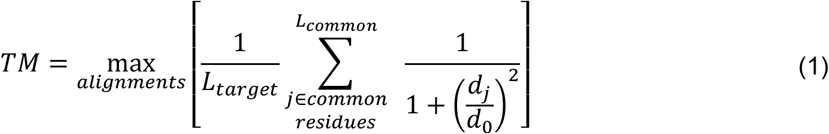

Each *d*_*j*_ is a distance between the predicted position of the Cα atom of residue *j* in the model and residue *j* in the experimental structure for a given superposition. A model of a protein can be superimposed in various ways on an experimental structure, and the maximum is taken over all possible alignments. In practice, the maximum is taken over only a subset of such alignments (e.g., by running different structure alignment programs or with different parameters). *d*_0_ is a scaling factor that reduces or eliminates the length dependence of the TM score for alignments of unrelated proteins (20). It has a fitted value of

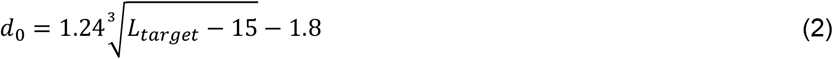

If *L*_*target*_ is 500 residues, *d*_0_ has a value of about 8 (Figure 1). The original TM score was used in development of protein structure prediction methods, when the sequence length of the experimental structure (the target) might be longer than the sequence length of the model (e.g., if only a single domain of the target could be modeled using templates). Thus, a partial model (or template) was heavily penalized. In some cases, the experimental structure might be missing some residues due to poor electron density. The sum is therefore over the number of residues the model and experimental structure have in common (*L*_*common*_).

**Figure 1.**
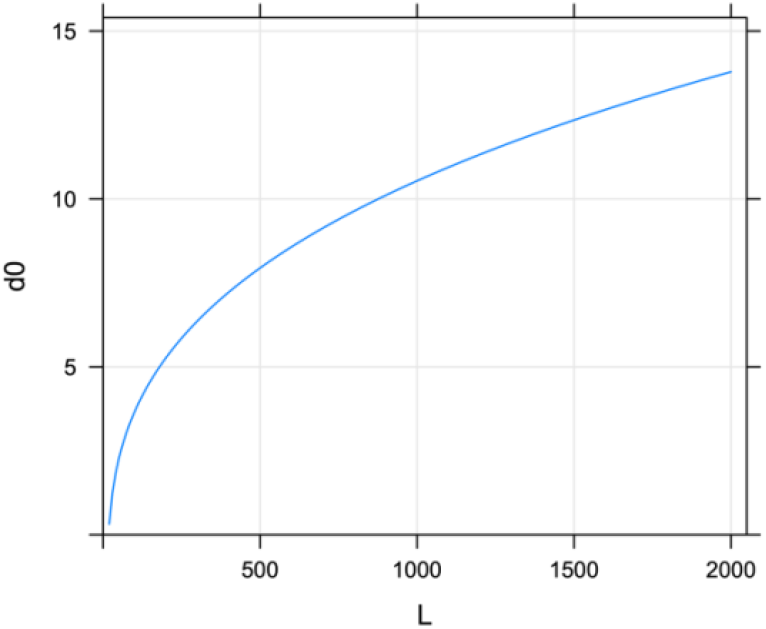
The *d*_0_ parameter in the TM score equation as a function of sequence length *L*.

AlphaFold2 and AlphaFold3 use the concept of “aligned error” to generate predicted accuracy metrics for output models (Figure 2). After superposing the N, Cα, and C atoms of residue *i* of a model onto the N, Cα, and C atoms of the same residue *i* in the experimental structure, the aligned error (AE) of residue *j* is the distance between the Cα atom of residue *j* in the model and Cα of residue *j* in the experimental structure. During training, the experimental structure is known, and the network is trained to predict a probability distribution over the aligned error distance when the experimental structure is not known (i.e., during inference). The probability distribution is defined over the *AE*_*ij*_ distance in 64 bins of width 0.5 Å (0Å-0.5Å, 0.5Å-1.0Å, …, 31.5-32Å), where the last bin also includes distances larger than 32 Å. The predicted aligned error, *PAE*_*ij*_, for each pair of residues is calculated from the predicted probability distribution over the aligned error with the equation (Eq. 11 in AF3 paper supplemental):

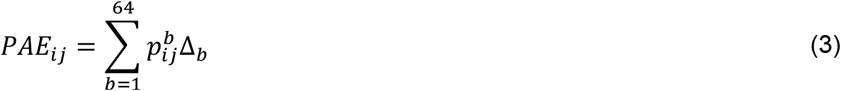

where Δ_*b*_ = (*b* − 0.5)⁄2 is the center of each bin (0.25Å, 0.75Å, …, 31.75Å), 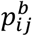 of bin *b*, and 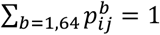.

**Figure 2.**
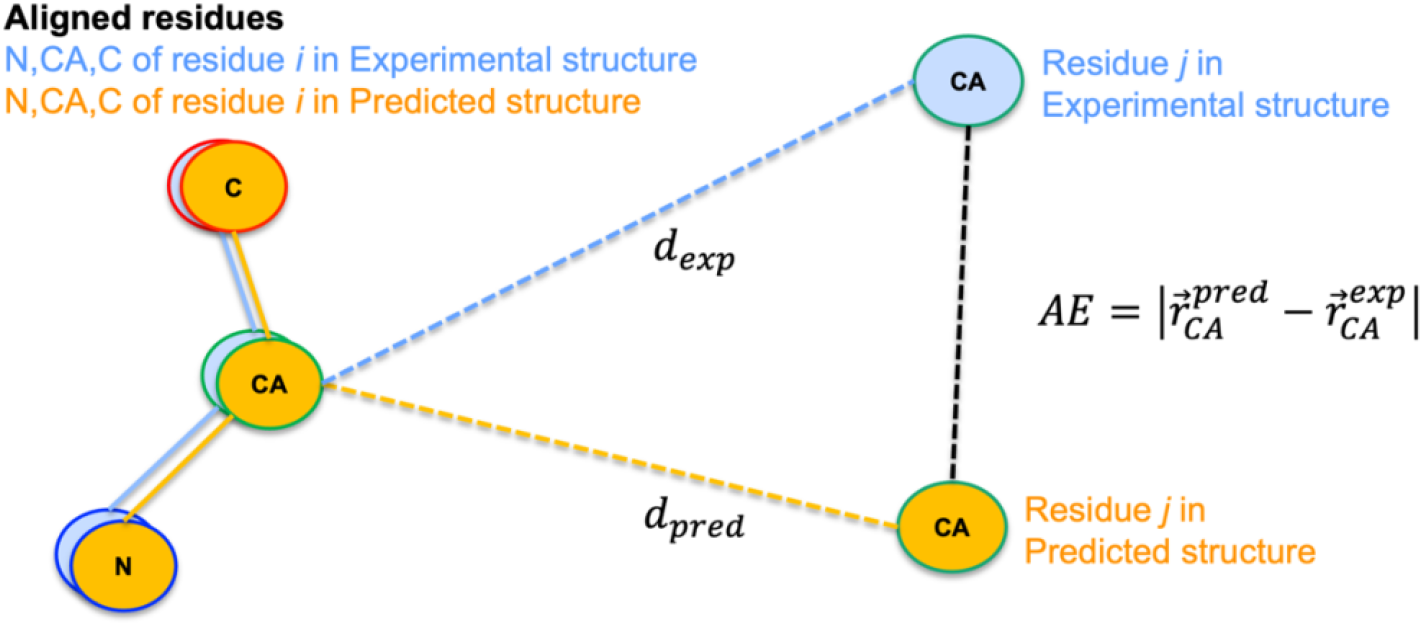
Definition of aligned error (AE) in AlphaFold2 and AlphaFold3.

For a single chain (or a whole protein complex), the *PAE*_*ij*_ values can be substituted into Equation 1 for the TM score to provide an equation for the *pTM* score (predicted template modeling score):

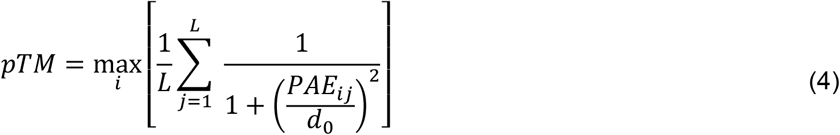

#### The role of residue *i* in this equation is to create a set of alignments used to calculate the TM score, one for each residue in the chain (or complex)

The value of *pTM* is then calculated from the highest scoring of these alignments, just as in Equation 1 for the original TM score

In the AlphaFold papers, the expression under the sum is instead calculated as an expectation value from the probability distribution of the aligned error used in Eq. 3 (AF3 paper supplemental Eq 17):

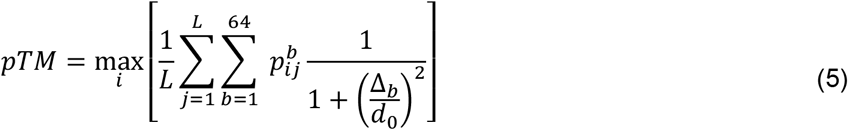

For simplicity, we define the pairwise *pTM* matrix from the aligned error probability distribution as:

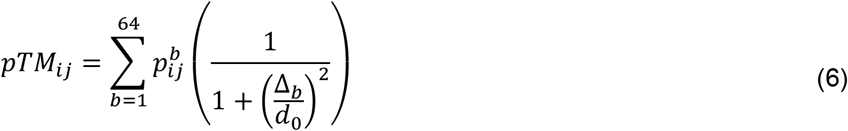

or alternatively (as an approximation) from the *PAE* value.

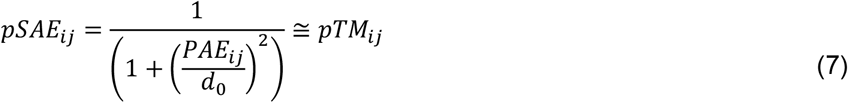

The *pSAE*_*ij*_ can be used anywhere *pTM*_*ij*_ can be used.

The residue-specific mean value of *pTM*_*ij*_, based on the alignment of residue *i* is given by:

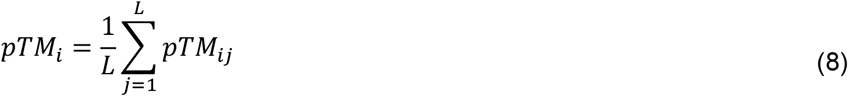

And 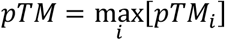. From these equations, we can generalize the expression for *pTM* by specifying the residue sets for the alignments (set *S*_1_, residues *i*) and those for the residue displacements between modeled structure and experimental structure, if it were known ( *S*_2_: residues *j*):

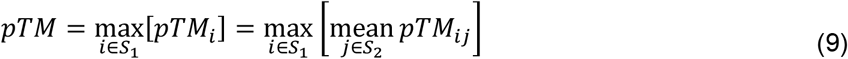

For a complex of two protein chains, A and B, we can perform the residue-residue structure superpositions over one chain (e.g., *S*_1_=chain A) and calculate the TM score over the other chain (*S*_2_=chain B), which would then contain a rotation-translation component as well as the accuracy of the structural model of chain B. So

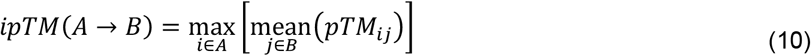

When AlphaFold2 or AlphaFold3 provides a value of *ipTM* for a pair of chains, it provides a single value which is the maximum of the two asymmetric values (or equivalently the maximum over all residues in both chains of the interchain *pTM*_*i*_):

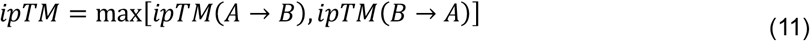

AlphaFold2 and AlphaFold3 use a value for *d*_0_ based on the sum of the lengths of the two chains.

AlphaFold3 provides an *ipTM* for each chain where the maximum is taken over all residues *i* in that chain and the mean is over all residues in all other chains.

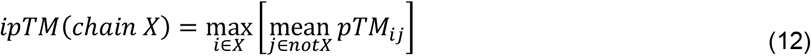

In AlphaFold2 and AlphaFold3, the overall *ipTM* of any multiprotein complex is calculated from the maximum over all residues of the mean *ipTM*_*ij*_, where the mean is taken over all residues in all other chains that do not contain residue *i*. The value of *d*_0_ is the sum of all protein chain lengths in the model.

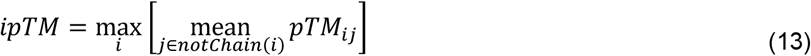

Our experience, and that of many others (14), demonstrates problems in the calculation of *ipTM* in the presence of disordered residues and other domains in the sequence constructs that do not interact between the chains. Frequently, users of AlphaFold2 and AlphaFold3 have to repeat calculations with different protein constructs that remove the disordered regions and observe a large change (up or down) in *ipTM*, even though the interacting domain-domain or domain-peptide complex structure remains the same. There are two scenarios to consider, one in which the ipTM score is artificially high and one in which it is artificially low. The reasons for both are clear from the equations presented above as follows.

If one chain is held fixed in sequence, while the other is extended to include disordered regions or additional domains that do not interact with the fixed sequence chain, the ipTM score ***goes up***. This is because the Δ_b_ values in Eq. 5 (or *PAE*_*ij*_ values in Eq. 7 if *PAE*s are used directly) are scaled down by the larger value of *d*_0_ (calculated from the sum of the two chain lengths, as in Eq. 2). This causes an artifact in the calculation of *ipTM*, artificially raising the score when the user adds sequence to one of the chains. Removing these regions in one chain lowers the score to more realistic values.

The opposite occurs when **both** chains contain non-interacting regions (disorder or non-interacting domains). For a protein-protein complex, *ipTM* is a mean value of *pTM*_*ij*_ over all residues *j* in one of the chains, after superposition on one residue *i* in the other chain (after taking the maximum over all residues *i*). It therefore includes many very low *pTM*_*ij*_ values because of the high *PAE*_*ij*_ values (>30) between ordered residues in one chain and disordered residues in the other. Any mobile domains that do not interact also lower the score, because they also contribute low *pTM*_*ij*_ from low *PAE*_*ij*_.

In practice, both effects are operative and compete unpredictably: raising *ipTM* due to an increase in sequence length and consequently *d*_0_ in Eq. 5, and lowering *ipTM* due to the inclusion of many poor PAE values for non-interacting sequence in both chains. As an example, we take the interaction between KRAS and the RAS-binding domain of RAF1 (Figure 3). When only the ordered domain sequences are input to AlphaFold-Multimer (v2.3), the *ipTM* is 0.85. When 120 residues of disorder are added to only one chain (the blue RAF1 in Example 2), the *ipTM* rises to 0.90 (since the ipTM is already quite high, it cannot raise significantly). This occurs because a residue in RAF1 (marked by a red asterisk) has high *pTM*_*ij*_ values with all residues in the fully ordered KRAS chain (magenta) and the value of *d*_0_ is higher in Example 2 than it is in Example 1. But when disorder is present in both chains (Examples 3 and 4 in Fig. 3), the *ipTM* is decreased because every residue in each chain has some low *pTM*_*ij*_ values with residues in the other chain. The decrease is proportional to the relative amount of disorder to order in the chain with *less* disorder. For example, with 120 disordered residues in each chain (Example 4), RAF1-RBD and KRAS are 61% and 41% disordered respectively. The *ipTM* value is due to residue T68 of RAF1, which sits in the interface with KRAS. Its pairwise *pTM*_*ij*_ values with KRAS residues are 59% ordered (at ∼0.9 each) and 41% disordered (at ∼0.2 each), or approximately 0.59^*^0.9 + 0.41^*^0.2 = 0.61 (the AF2 output value is 0.56).

**Figure 3.**
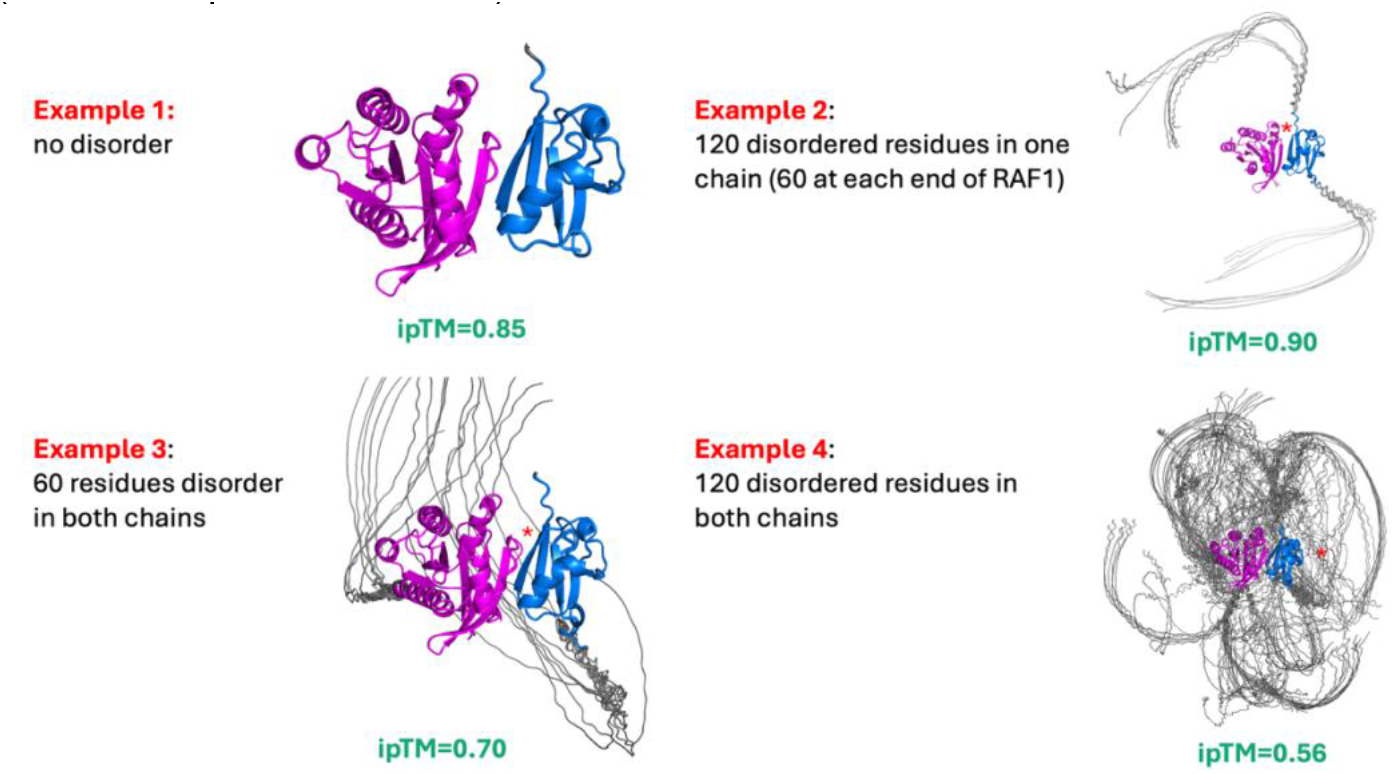
AlphaFold2 models of the complex of KRAS (magenta) and the RAS-binding domain of RAF1 (blue). Disordered residues (15 repeats of the sequence GGGS) were added to the N or C terminus (or both) of one or both chains, which increases the *ipTM* when this occurs in one chain (Example 2) and decreases it when it occurs in both chains (Examples 3 and 4).

There are several ways of dealing with this. In the *ipTM* expressions, we could skip residue pairs where one (or both) residues have *pLDDT* values less than some cutoff value (e.g. *pLDDT*<50). This does not always work: auxiliary domains in one or both of the proteins that do not contribute to the protein-protein interaction will have good *pLDDT*s but poor intermolecular *pTM*_*ij*_, thus lowering the *ipTM*.

The *ipTM* could be calculated over only contacting residues in the model within some cutoff distance. This can also be a problem because disordered residues or auxiliary domains in one or both chains can contact the other chain and contribute poor *pTM*_*ij*_ to the *ipTM* evaluation,

Varga et al recently proposed using the predicted distance distograms produced by the AlphaFold2 network to restrict the calculation of *ipTM* to interchain residue pairs that are predicted to be in contact (21). This method excludes disordered regions and auxiliary domains that do not have a strongly predicted interaction, even if they are in contact in the models. Their score, called *actifpTM*, is calculated over the subsets of residues that make up the interface of two chains but only those that AlphaFold2 is confident about. *actifpTM* is now implemented within the ColabFold framework (22).

We propose another alternative, where we use the *PAE*_*ij*_ values to restrict the *ipTM* calculation to interchain residue pairs that have well predicted aligned error distances, regardless of whether they are in or near the protein-protein interface. In contrast to *actifpTM*, we adjust the value of *d*_0_ in the asymmetric *ipTM* expression to the number of residues in the chain under the mean expression with good interchain *PAE* values (Equation 10). This is critical, because a small number of interchain residue pairs with spuriously good *PAE*_*ij*_ and consequently good *pTM*_*ij*_ values may produce an unrealistically *ipTM* if *d*_0_ is not adjusted.

We define the *ipSAE* score (*i*nterface *p*redicted TM *S*core based on *A*ligned Errors) for two chains (A and B) as follows:

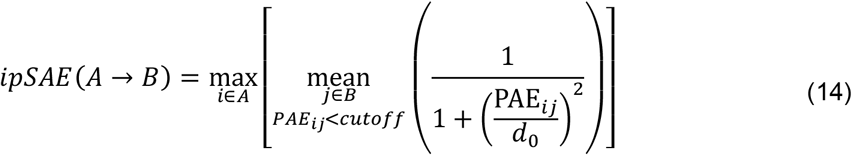

And

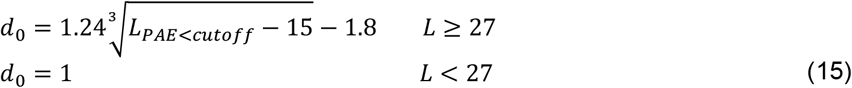

Here, *L*_*PAE*<*cutoff*_ is the number of unique residues in chain B that have *PAE*_*ij*_ < *cutoff* given the identity of the aligned residue *i*. We use a minimum value of 1 for *d*_0_, since Yang and Skolnick did not test the fit for proteins shorter than 30 amino acids (*d*_0_=1 for L∼26.5), and the denominator in Eq. 14 starts to blow up for values << 1.0, which may not be realistic or helpful. In the AlphaFold code, the minimum value is set to 19, since *L*=18 produces a negative number.

For a given chain pair, the *ipSAE* score is the maximum of the two asymmetric values:

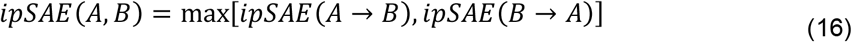

*ipSAE* can be calculated for every pair of chains in a multi-chain complex from the *PAE* matrices from the AlphaFold2 or AlphaFold3 output json files.

## Results

### RAF1 complexes

The TKL family kinase, RAF1, contains three domains: a RAS-binding domain (RBD: residues 56-131), an immediately adjacent cysteine-rich domain (CRD: residues 138-184), and a protein kinase domain (PK: residues 340-614). The rest of the chain of length 648 residues is intrinsically disordered (residues 1-55, 185-339, and 615-648).

As noted above, AlphaFold-Multimer models of the RAF1-RBD with KRAS result in *ipTM* values that are lowered in the presence of artificial disordered sequences when they are present in both chains (Figure 3). The *ipSAE* value of 0.8 for Example 4 eliminates most of the disorder effect.

RAF1 interacts with the TKL family pseudokinase KSR1, facilitating the activation of RAF1 and its translocation to the membrane (23). KSR1 also contains three domains: the coiled-coil/Sterile-α-motif domain (CC-SAM: residues 30-172), a cysteine-rich domain, homologous to the CRD of RAF1 (CRD: residues 347-391), and a protein pseudokinase domain (pPK: residues 599-833). There is no experimental structure of a RAF1-KSR1 complex, but it has been hypothesized that the two kinase domains bind in a mode similar to the well-known BRAF homodimer (24).

AlphaFold-Multimer models of the heterodimer sequences of full-length RAF1 and full-length KSR1 show a kinase/pseudokinase heterodimer that is very similar to the BRAF homodimer (24) with *ipTM* values between 0.38 and 0.41 across 25 models (5 seeds by all 5 sets of AF2 model weights, without templates). The other folded domains and disordered regions of both proteins are not in fixed position relative to the kinase domains across the 25 models, and do not show any key interprotein interactions (Figure 4A).

**Figure 4.**
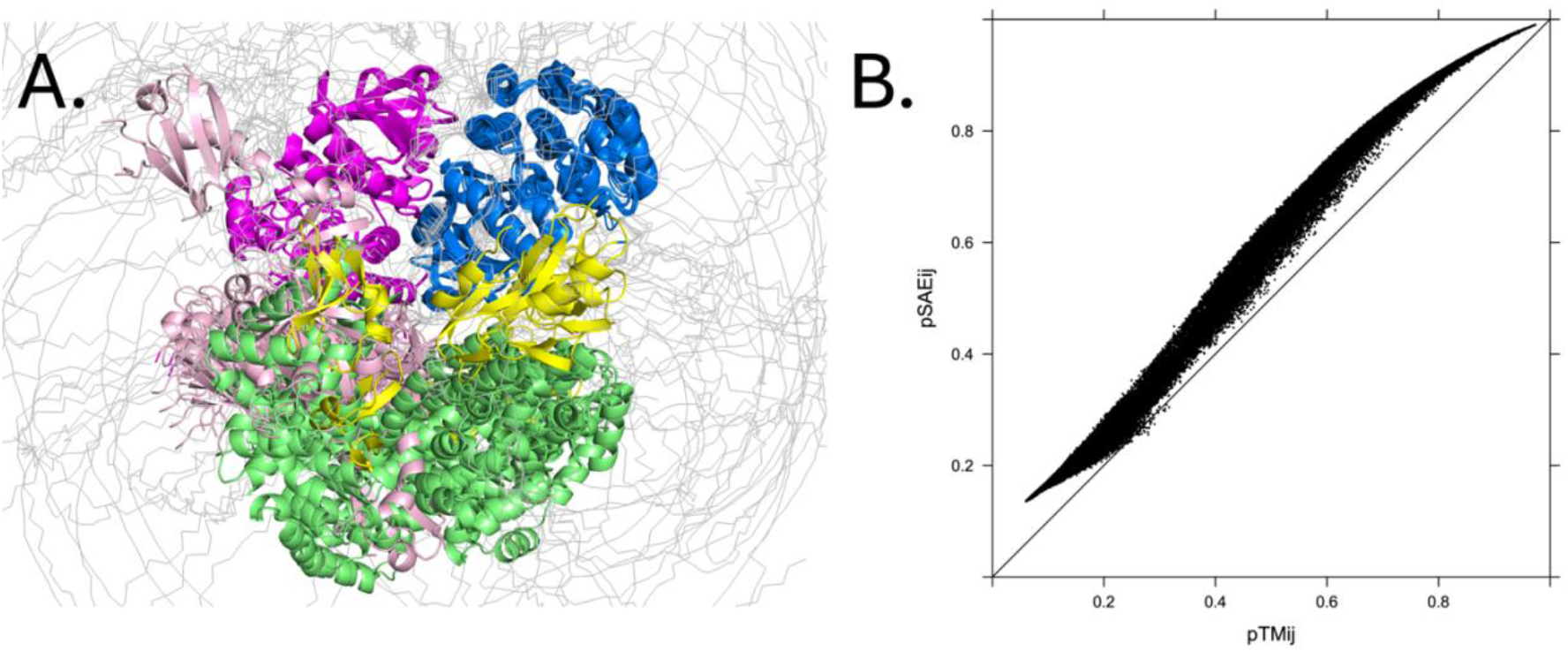
RAF1-KSR1 models. **A. AlphaFold2 models of full-length RAF1 and KSR1**. RAF1 RBD-CRD domains in pink and kinase domain in magenta. KSR1 CC-SAM domains in green, CRD in yellow, and pseudokinase domain in blue. The 25 models are aligned on the kinase domain of RAF1. The non-kinase domains are not fixed relative to the two kinase domains. **B. Scatterplot of interchain *pSAE***_**ij**_ **vs *pTM***_**ij**_ **values for RAF1-KSR1 complex**. *pTM*_*ij*_ was calculated from the *pTM* matrix output by AlphaFold2 (from a modified version of ColabFold). It uses a *d*_0_ from the combined length of both proteins. *pSAE*_*ij*_ was calculated with no *PAE* cutoff and with a *d*_0_ also based on the combined length of both protein chains.

As described above, we can use the *PAE* values to calculate *ipTM*-like scores over specified interprotein residue pairs. AF2 calculates the full *pTM* matrix with a value of *d*_0_ that is the combined length of the two proteins. If we calculate *pTM* with the *PAE* values instead and use the same *d*_0_, the interchain *pSAE*_*ij*_ and *pTM*_*ij*_ are highly correlated (Figure 4B).

AlphaFold-Multimer calculates the *ipTM* via Equations 10 and 11 by calculating the *pTM*_*i*_ for each residue in both chains (Figure 5). The two kinase domains are responsible for the high scoring regions, while the accessory domains in each protein are visible as small bumps in the plots. The maximum value in the curve in Figure 5 occurs for residue W632 of KSR1, which is in the interface between the kinase domains. If we use the *pSAE*_*ij*_ value to calculate *pSAE*_*i*_ for each aligned residue *i* of the complex and adjust *d*_0_ for the number of residues that have a good *PAE* for the aligned, we see higher scores for the kinase domains in RAF1 and KSR1 and zero for the accessory domains. AlphaFold2’s *ipTM* for the top-ranking complex is 0.41, while the *ipSAE* score is 0.73.

**Figure 5.**
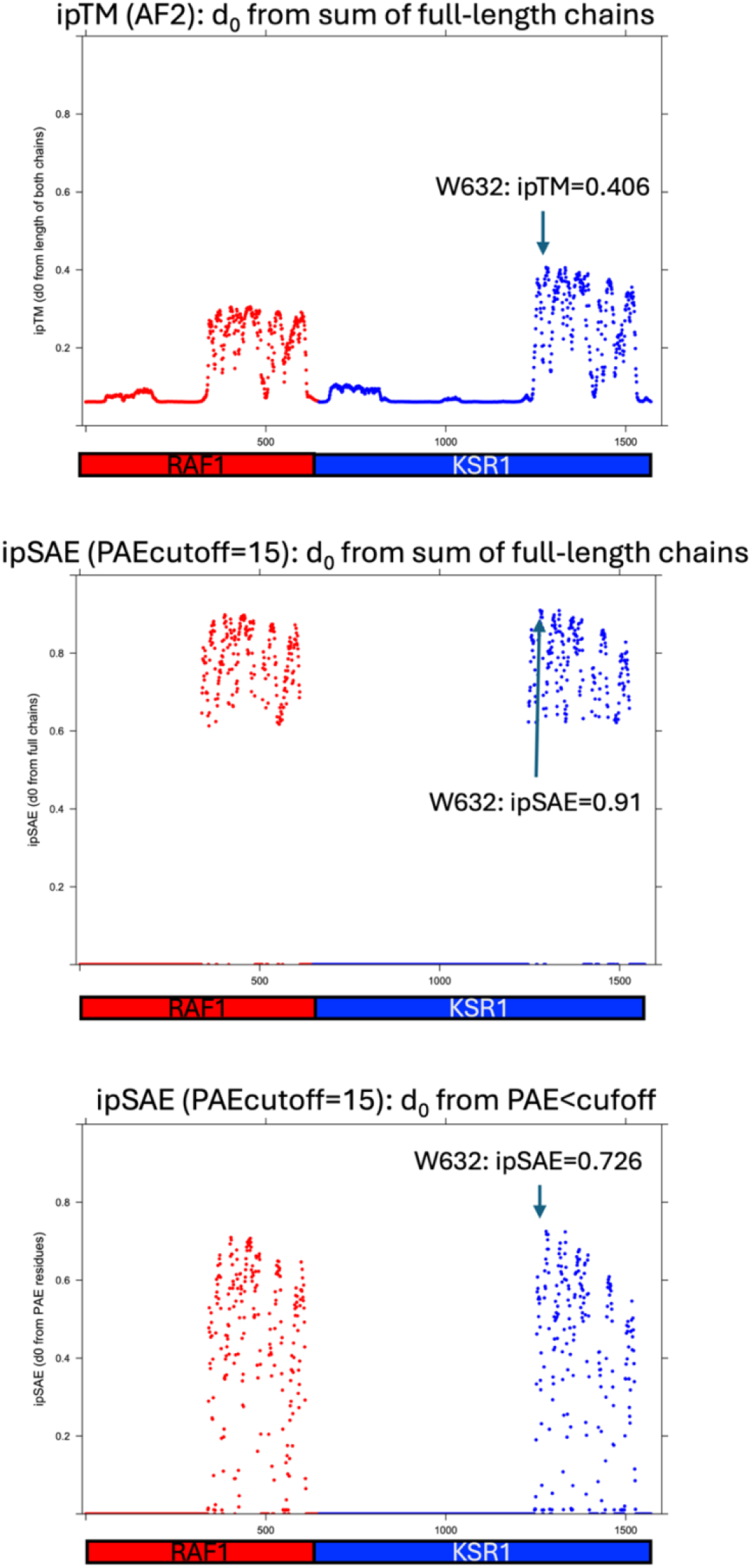
Per-residue *ipTM* and *ipSAE* scores for the RAF1-KSR1 complex. *ipTM* (top) was calculated from the pairwise *pTM* matrix from AlphaFold-Multimer v2.3 (ColabFold). It uses a *d*_0_ from the combined length of the two proteins (1571 amino acids; *d*_0_=12.57). *ipSAE* with a *PAE* cutoff of 15.0 and the same value of *d*_0_ (middle figure). A cutoff for the *PAE* score of 15.0 Å was used to produce the *ipSAE* scores (bottom figure). The maximum occurs for W632 of KSR1 for both scores (green arrows) with a value of *d*_0_ of 6.32 (286 residues in RAF1 with *PAE*<15 Å).

The TKL family kinase, RIPK1, is not known to bind to RAF1. RIPK1 has a kinase domain (PK: residues 8-324), a RIP homotypic interaction motif (RHIM: residues 531-547), and a Death domain (DD: residues 567-671). *ipTM* and *ipSAE* plots by residue are shown in Figure 6, demonstrating the effects of using the *PAE* cutoff and the evaluation of *d*_0_ based on the number of residues with *PAE* less than a cutoff of 15 Å. The top plot shows the per-residue *ipTM* scores from AlphaFold2 (by averaging each row of the interchain *pTM*_*ij*_ values output from a modified version of ColabFold). AF2 uses a *d*_0_ based on the sum of the two chain lengths (in this case 648 + 671 = 1319 residues, *d*_0_=11.75). In the middle plot, the *PAE* matrix is used to limit the number of *pTM*_*ij*_ used for each residue. It uses the same value of *d*_0_ (11.75) as in the top plot. The resulting residue-specific *ipSAE* values are much higher than the *ipTM* values, with an overall *ipSAE* value of 0.459 from the alignment on residue L433. This is expected because residue pairs with good *PAE* values will have high pairwise *ipTM* (or *ipSAE*) values. But the number of such pairs in truly non-interacting proteins is quite low, if AlphaFold is working as expected. In the bottom plot, the combined effect of the *PAE* cutoff of 15 Å and the residue-specific *d*_0_ values bring the residue *ipSAE* values way down. The overall value of *ipSAE* is now 0.044, indicating that the proteins are not likely to interact. The value of *d*_0_ was 3.05 from 75 residues below the *PAE* cutoff.

**Figure 6.**
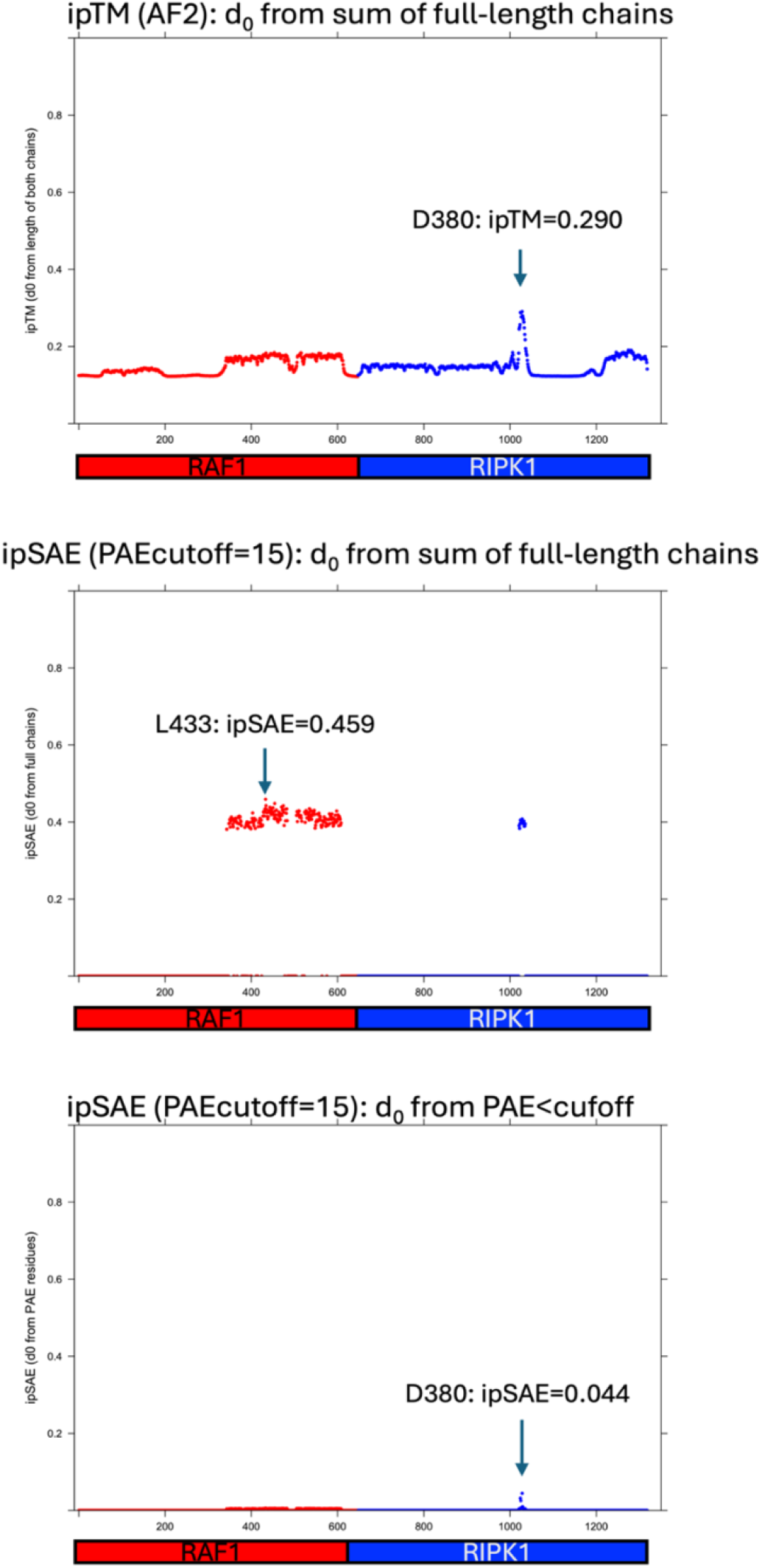
Per-residue *ipTM* (top) and *ipSAE* scores (middle and bottom) for the RAF1-RIPK1 pseudocomplex. A cutoff for the *PAE* score of 15.0 Å was used to produce the *ipSAE* scores. The *ipTM* score per residue scores (top plot) show a modest interaction between the chains with a maximum value at residue D380 of RIPK1, *ipTM*=0.290. AF2 uses *d*_0_ from the sum of the full chain lengths. A *PAE* cutoff using the same *d*_0_ (middle plot) (from the sum of both chain lengths) raises the *ipSAE* values compared to the *ipTM* values. But adjusting *d*_0_ to account for the number of residues in the mean *pTM*_*ij*_ calculation (e.g. for each residue in RAF1, this is the number of residues in RIPK1 that have *PAE*<*PAEcutoff*) significantly lowers the score of the non-interacting proteins to a value of 0.044.

The TKL family kinase, RIPK1, is not known to bind to RAF1. RIPK1 has a kinase domain (PK: residues 8-324), a RIP homotypic interaction motif (RHIM: residues 531-547), and a Death domain (DD: residues 567-671). *ipTM* and *ipSAE* plots by residue are shown in Figure 6, demonstrating the effects of using the *PAE* cutoff and the evaluation of *d*_0_ based on the number of residues with *PAE* less than a cutoff of 15 Å. The top plot shows the per-residue *ipTM* scores from AlphaFold2 (by averaging each row of the interchain *pTM*_*ij*_ values output from a modified version of ColabFold). AF2 uses a *d*_0_ based on the sum of the two chain lengths (in this case 648 + 671 = 1319 residues, *d*_0_=11.75). In the middle plot, the *PAE* matrix is used to limit the number of *pTM*_*ij*_ used for each residue. It uses the same value of *d*_0_ (11.75) as in the top plot. The resulting residue-specific *ipSAE* values are much higher than the *ipTM* values, with an overall *ipSAE* value of 0.459 from the alignment on residue L433. This is expected because residue pairs with good *PAE* values will have high pairwise *ipTM* (or *ipSAE*) values. But the number of such pairs in truly non-interacting proteins is quite low, if AlphaFold is working as expected. In the bottom plot, the combined effect of the *PAE* cutoff of 15 Å and the residue-specific *d*_0_ values bring the residue *ipSAE* values way down. The overall value of *ipSAE* is now 0.044, indicating that the proteins are not likely to interact. The value of *d*_0_ was 3.05 from 75 residues below the *PAE* cutoff.

### Benchmark of recent PDB entries

We identified a set of 40 PDB entries that share at most 40% identity with any chain present in the PDB prior to the AlphaFold-Multimer v2.3 cutoff date of Sept. 30, 2021. The entries had to have exactly two unique sequences and have a biological assembly consistent with a pairwise interaction of the two unique sequences (e.g., we excluded assemblies larger than octamers and chose entries where the shorter sequence interacted with only one copy of the longer sequence). Each sequence had to have at least 12 amino acids in the coordinates of the PDB file. Sequence identities were obtained from the PISCES webserver (25). We ran AlphaFold-Multimer v2.3 on the PDB sequences themselves and from the full-length Uniprot sequences, as identified from SIFTS (26) (as given in the PISCES sequence files). We also created a set of 70 AlphaFold jobs by randomly creating heterodimer pairs by mixing sequences from different entries in the set of 40 PDB entries. These were run with the full-length Uniprot sequences only.

The results of the *ipTM* and *ipSAE* scores are shown in Figure 7. The top left panel shows the *ipTM* values calculated from the *pTM* matrix in ColabFold. AF2 uses a value of *d*_0_ calculated from the sum of the lengths of the two sequences in each query. Our values of *ipTM* agree exactly with the *ipTM* values present in the AF2 json output files. If we use the *PAE* values in the *ipTM* expression (but no *PAE* cutoff), instead of the AF2 *pTM* matrix, we get quite similar distributions (top right panel). In both panels, there is overlap in the density between values of *ipTM* or *ipSAE* from 0.3 to 0.7 for the true dimers (full-length Uniprot sequences, blue curves and data points) and false dimers (full-length Uniprot sequences, magenta curves and data points).

**Figure 7.**
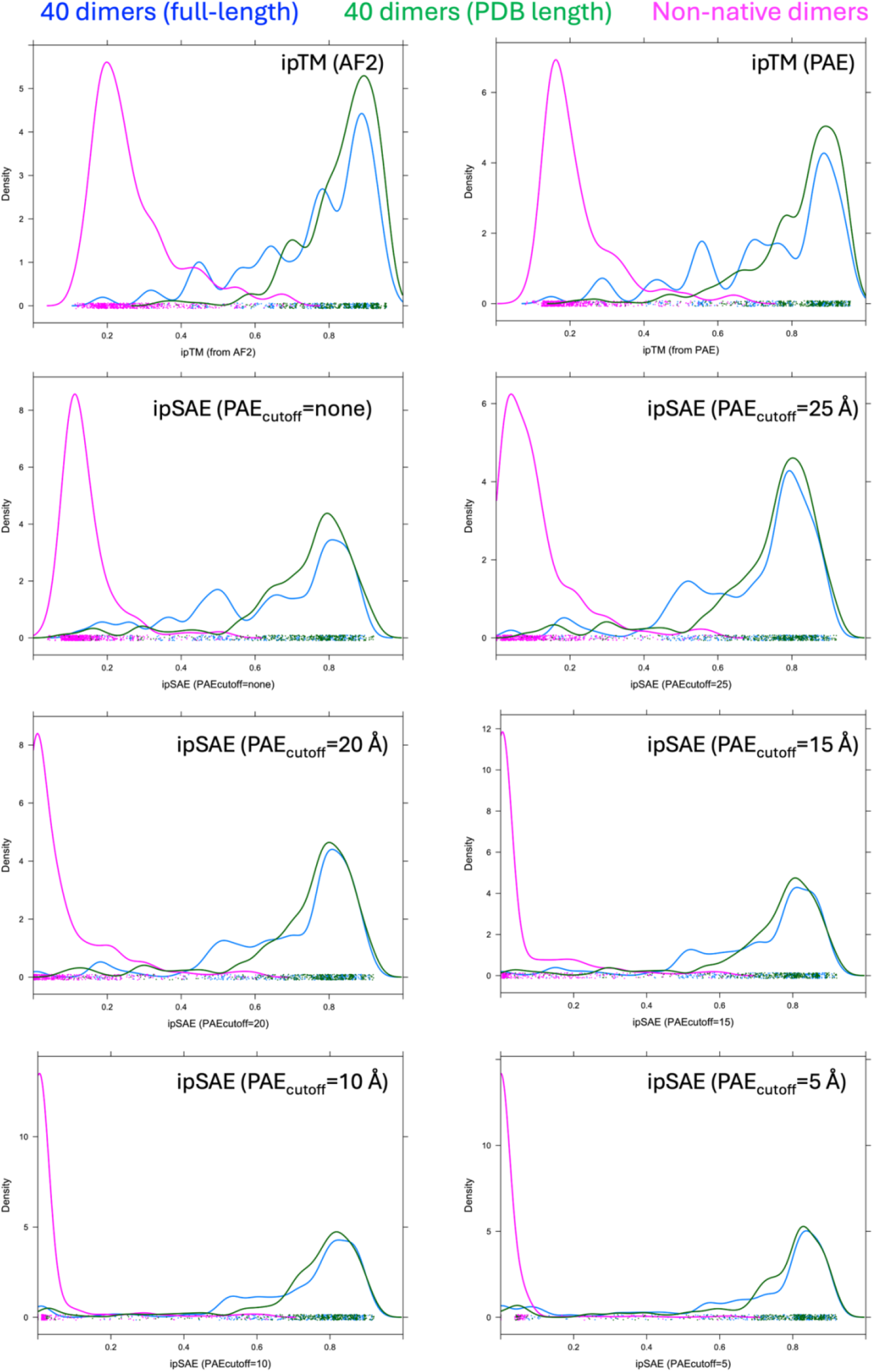
Benchmark of recent PDB heterodimers. A set of 40 heterodimer PDB entries with less than 40% sequence identity to any sequences in the PDB prior to October 1, 2021 were identified. The PDB sequences were used as queries to AlphaFold-Multimer v2.3 (green curves). The full-length Uniprot sequences for these chains were also used as a second set of 40 target complexes for AF-Multimer v2.3 (blue curves). A third set of 70 targets was built from mixing the Uniprot sequences from different entries (magenta curves). The plots show kernel density estimates of *ipTM* and *ipSAE* for the top 10 ranked complexes (AF2 ranking based on 0.8^*^*ipTM* + 0.2^*^*pTM*) out of 25 models (5 seeds x 5 AF2 weight-sets with no templates used). The top left panel shows *ipTM* based on the *pTM* matrix from AF2. The top right panel shows *ipTM* calculated from the *PAE* matrix instead of the *pTM* matrix (the *PAE* value is used in the denominator of the *pTM* expression instead of the sum over probabilities). The remaining rows show *ipSAE* values with different *PAE* cutoffs used in the mean value calculation. *d*_0_ for these calculations was based on the *PAE* cutoff. The set of 40 PDB entries is: 7f4p, 7qii, 7sck, 7t5p, 7tj4, 7wmv, 7wwq, 7ytu, 7zd5, 8a51, 8a82, 8bfj, 8blw, 8cdp, 8dqv, 8fbd, 8fzz, 8g0p, 8gs1, 8guo, 8hi7, 8hk0, 8ir4, 8jj9, 8jmq, 8jzd, 8orn, 8ows, 8q4h, 8qvc, 8r5i, 8s2m, 8vjl, 8vx9, 8wx5, 8xfb, 8y2n, 8ypu, 8zlz, 9dk1.

**Figure 8.**
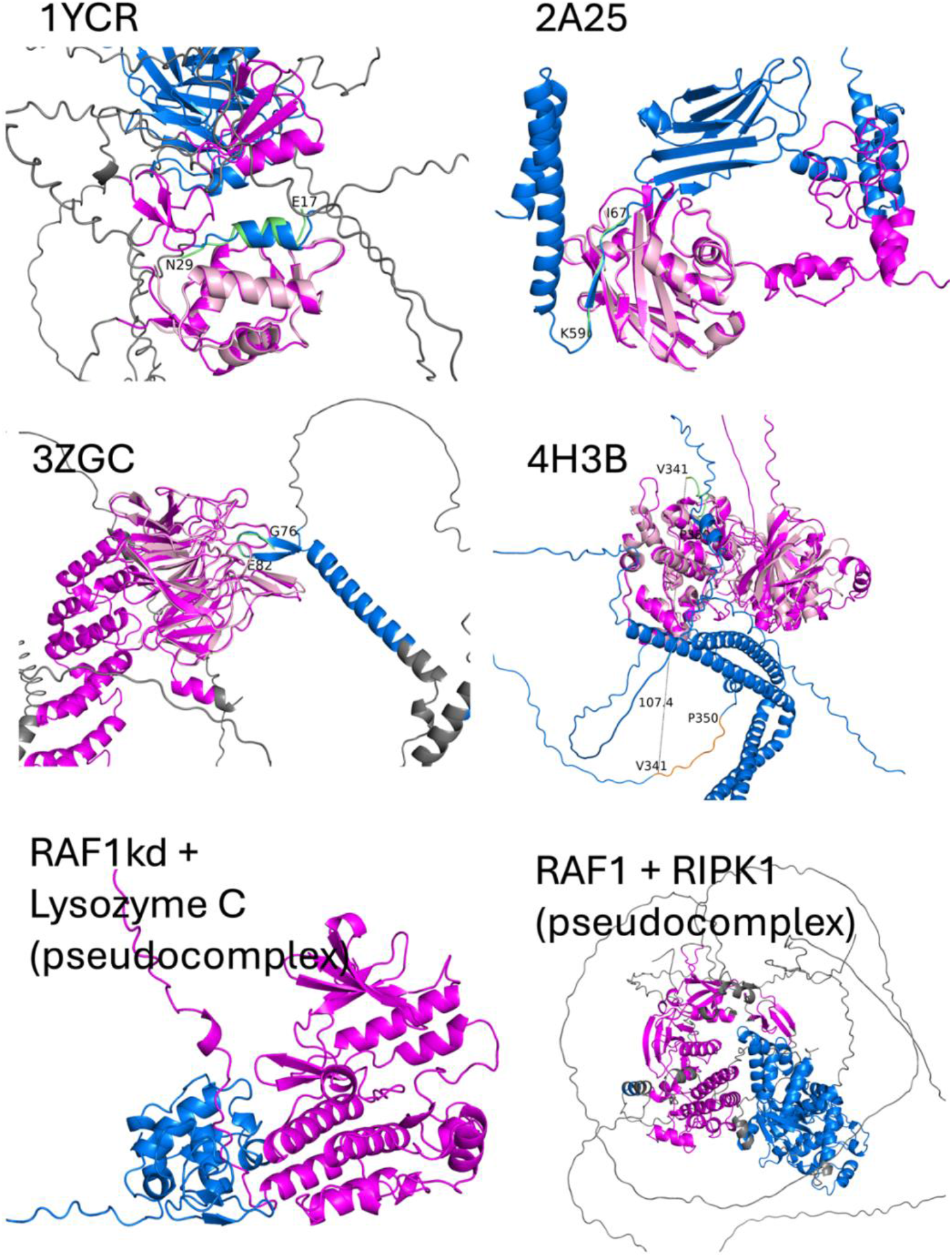
Top-ranked ColabFold AlphaFold-Multimer v2.3 models of protein complexes from full-length Uniprot sequences. PDB entries 1YCR, 2A25, 3ZGC, and 4H3B were used as examples in the preprint of Varga et al. RAF1 kinase domain plus chicken Lysozyme C and full-length RAF1 with RIPK1 are examples of non-interacting proteins and their resulting scores.

In the next three rows, kernel density plots of *ipSAE* values are shown for all three sets of targets with different values of the *PAE* cutoff in descending order (32 = no cutoff, 25, 20, 15, 10, and 5 Å). As the *PAE* cutoff decreases, the density in the mid-range of *ipSAE* decreases, separating true from false dimers more effectively than the *ipTM* values from AlphaFold (top left panel). The true dimers with Uniprot sequences have significantly improved *ipSAE* values at lower cutoffs, because they contain disorder and accessory domains that do not form part of the interaction between the two proteins. The PDB sequences (green curves), conversely, do not change that much with the *PAE* cutoff since they do not usually contain disordered regions or mobile domains that do not form part of the interaction. The overall results indicate that the *ipSAE* score may be better at separating true from false interactions even in the presence of disordered sequences and/or accessory domains in both sequences. **Cutoffs of 10 or 15 Å may be most suitable**.

### Comparison with actifpTM

Varga et al. (21) identified the same problem with the *ipTM* score as we have discussed above – that disordered regions depress the *ipTM* score when they are not part of the binding interface between two proteins. They gave four example systems of protein-peptide complexes: PDB entries 1ycr (MDM2 and P53 peptide), 2a25 (E3 ubiquitin ligase SIAH1 and Calcyclin binding protein peptide), 3zgc (KEAP1 and NF2L2 peptide), and 4h3b (MAPK10 and SH3 domain-binding protein 5 peptide). We ran AlphaFold-Multimer v2.3 on the Colabfold Jupyter notebook, which calculates the *actifpTM* values, using the full-length Uniprot sequences of both chains (instead of the PDB constructs or short elongations of these, used in the *actifpTM* preprint). The calculations were performed with two seeds, no templates, and 3 recycles. We calculated *ipSAE* at different *PAE* cutoffs on the rank001 models from Colabfold. The results are shown in Table 1. For three of the targets, AlphaFold produces good models where the binding peptide is correctly placed on the folded domain, even though the full-length Uniprot sequence was provided to Colabfold. After superposition onto the folded domain from the PDB structure (chain A in all cases), the RMSDs were 1.32, 0.72, and 1.12 Å for entries 1ycr, 2a25, and 3zgc. For these three entries, the *actifpTM* values were quite high, ranging from 0.93 to 0.97. The *ipSAE* values were lower with values around 0.68, 0.55, and 0.73 Å respectively (at *PAE* cutoff 10 Å).

**Table 1.**
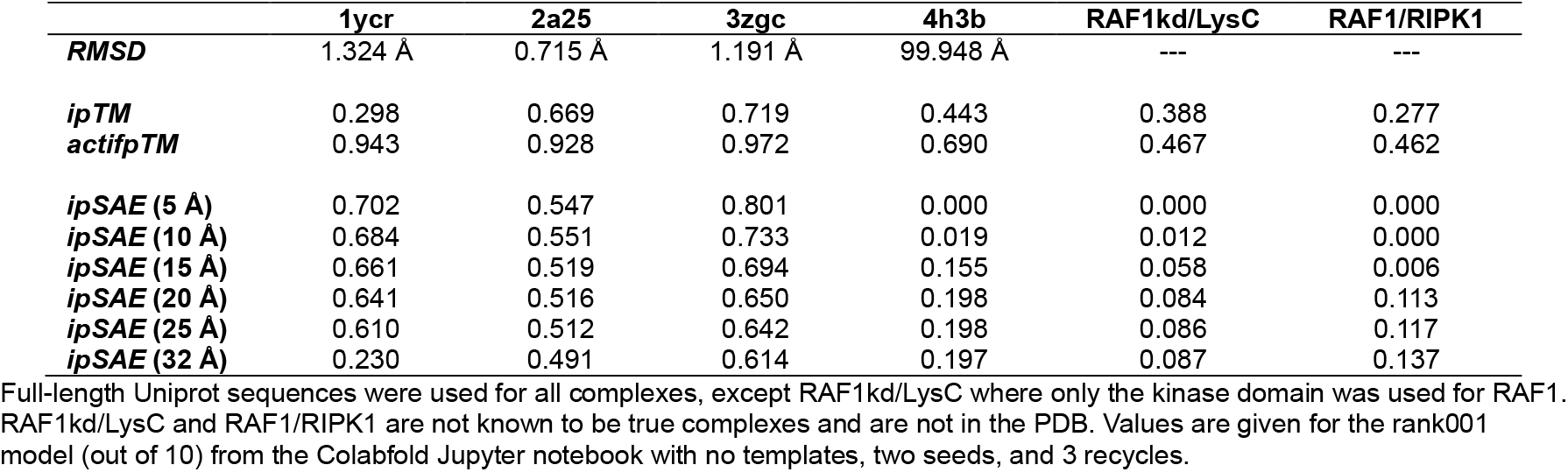
Comparison of *actifpTM* targets with their *ipTM* and *ipSAE* values.

The 4h3b structure is quite different. AlphaFold places the wrong peptide from SH3BP5 into the inhibitory binding site on the kinase domain MAPK10. In the PDB structure, residues 341-350 bind to the kinase domain. But in the model from full-length Uniprot sequences, the 341-350 segment is 100 Å away. Instead, residues 425-439 bind to the kinase domain in the SH3BP5 binding site. The *ipTM* from AlphaFold is 0.443, and the *actifpTM* value is 0.690, while the *ipSAE* values range from 0.0 (no *PAE* pairs less then 5 Å) to 0.20 (*PAE* cutoff 25 Å).

We also ran Colabfold calculations of the RAF1 kinase domain with a presumably non-interacting protein, chicken lysozyme C (LYSC_HUMAN), and full-length RAF1 with RIPK1. For LYSC, The *ipTM* value was 0.388 and the *actifpTM* was 0.467. The *ipSAE* scores successfully identify the non-interaction with *ipSAE* values from 0.0 to 0.1 (Table 1, last column). For RIPK1, *ipTM* was 0.277, *actifpTM* was 0.462, and *ipSAE* was 0.0 (*PAE* cutoffs ≤ 15 Å).

### Structure prediction benchmark of Genz et al

Genz et al recently published a benchmark of 223 heterodimeric complexes absent from the training data of AlphaFold2 and AlphaFold3 (6). They ran ColabFold with and without templates (labeled CF-T and CF-F respectively) and AlphaFold3 (AF3) and assessed the correlation of various metrics with structural accuracy as measured by DockQ. The sequences used in these computations were those from the constructs in the PDB structures. While the main advantage of the *ipSAE* score may be in handling sequences with high amounts of disorder or non-interacting domains in one or both proteins, we wanted to assess whether *ipSAE* had good correlation with structure prediction accuracy. Luca Genz was kind enough to run their benchmark with the *ipSAE* score and compare it to the other metrics in their published paper. The results are shown in Fig. 9. *ipSAE* performs slightly better than the next best metrics on the ColabFold models and equivalently on the AF3 models.

**Figure 9.**
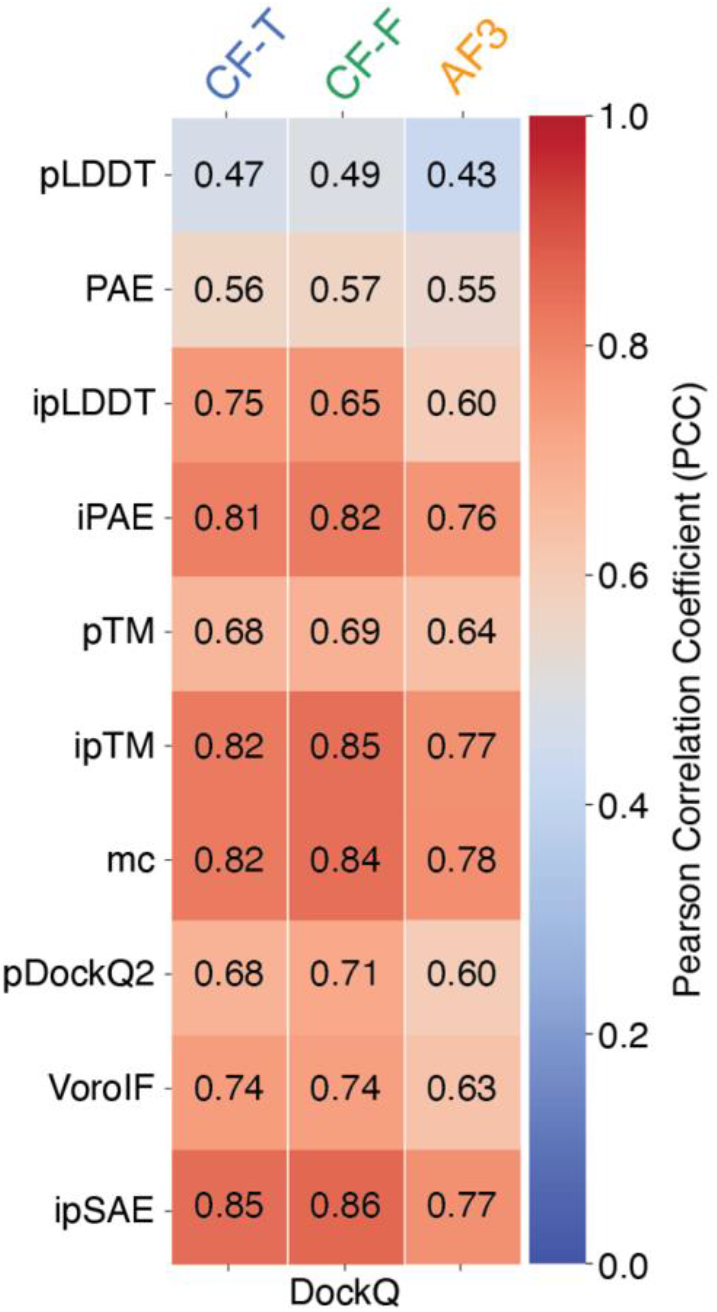
Benchmark of Genz et al. for correlation of various scoring metrics with structure prediction accuracy as measured by DockQ. The set of targets comprised 223 heterodimers from the PDB not in the training data of AlphaFold2 and AlphaFold3. CF-T and CF-F indicate template-based and template-free modeling with ColabFold.

## Discussion

We have proposed an *ipTM*-like score based on the output of AlphaFold2. The code has been modified to work on AlphaFold3 output, both from the AlphaFold3 server and AlphaFold3 run locally, as well as Boltz1. It works on modified amino acids as well as protein/nucleic acid interactions.

The *ipSAE* score is calculated over interchain residue pairs that pass a *PAE* cutoff, thus eliminating the effect of disordered regions in both chains and/or accessory domains that AlphaFold2 does not predict to be part of the binding interface. On a benchmark of 40 heterodimer complexes in the PDB not very similar (at 40% sequence identity) in the AlphaFold2 training set and 70 non-interacting sequence pairs from the same set, models based on full-length Uniprot sequences showed greater discrimination between true and false dimers with the *ipSAE* score compared to *ipTM*. We also showed that in some cases, our score behaves better at discrimination true than false interactions than the recently proposed *actifpTM* score of Varga et al. A true comparison would require a much larger set of targets.

Like *ipTM* in AlphaFold3 output and the *actifpTM* score, the *ipSAE* score can be calculated for every pair of chains in a multi-chain complex from AlphaFold2 output. We calculate the asymmetric values (A⟶B is different from B⟶A, where the first chain contains the aligned residues and the second chain contains the scored residues in the *PAE* values), as well as the maximum over all residues in both chains. It is possible there is insight to be gained in considering both values, rather than just the maximum, particularly for protein-peptide complexes.

Further comparison is needed to other scores presented in the literature that account for the flaws in *ipTM* in various ways. We made a few comparisons to the *actifpTM* score (21), which like *ipSAE* limits (and weights) the contribution of pairwise *ipTM* matrix elements to the resulting score. Kim et al. presented to the Local Interaction Score (27), which is obtained from the by converting *PAE* scores to a score from 0 to 1.0 and averaging over all interchain residue pairs with *PAE* ≤ 12 Å. The pDockQ (28) and pDockQ2 (18) scores are based on the *pLDDT* and *PAE* scores of interface residues, and also attempt to improve on the *ipTM* score from AlphaFold.

In addition to its ability to distinguish true from false interacting pairs, even in the presence of substantial disorder and non-interacting accessory domains, a recent benchmark from Genz et al. demonstrate that ipSAE performs as well or slightly better than *ipTM* as demonstrated by correlation with DockQ scores.

The *d*_0_ parameter in the TM expressions presents challenges for short peptides. In the original TM score paper, no individual structures were compared that were shorter than 40 amino acids. *d*_0_ becomes negative when the protein length is less than 19 amino acids, with a resulting *d*_0_ value of 0.17 when the length is 19. But in that case, the denominator of the *pTM* expression blows up and becomes very low, not matter how accurately the position of a peptide bound to a folded domain is predicted. To avoid this, we chose to set a minimum value of *d*_0_ to 1.0, which is a peptide length of approximately 27 amino acids. But this is somewhat arbitrary and needs to be investigated further.

Finally, the method for calculating the *ipTM* in the AlphaFold programs relies on the maximum *ipTM* over the residues in both chains. But many protein pairs have multiple domain-domain interactions separated by disordered regions. In these cases, the *ipTM* only scores one domain-domain pair (which ever scores highest) and the other(s) do not contribute. Examination of the *PAE* plot is helpful in identifying such cases. Models can then be produced with shorter constructs to estimate the *ipTM*s of each domain-domain interaction. Outputting *ipSAE* values from different aligned residues (not just the maximum value) may be useful in deriving a more useful metric than the methods described here and elsewhere. Our script, described below, outputs a file with the by-residue values of *ipSAE* which may be used for this purpose.

Recently, several papers have utilized the *ipSAE* score in protein binder design and structure prediction of protein complexes. Overath et al assembled a set of more than 3,700 designed binders with experimental data, comprising 436 binders and 3,324 non-binders, from studies on 15 different targets from various labs (29). The minimum of the two asymmetric AlphaFold3 *ipSAE* scores (target⟶binder and binder⟶target) was found to have 1.4 times the precision of *iPAE* used for binder design in the RFDiffusion program (30). It was the single best predictor of binding compared to *ipTM, iPAE, actifpTM*, and *pDock*Q. Chow et al. (31) designed 1,589 novel protein binders targeting BCMA, CD19, and CD22 with BindCraft (32) and RFDiffusion. They found that *ipSAE* was a better metric for identifying binders than *ipTM* and *iPAE*, and correlated well with kD. Moriwaki et al found that *ipSAE* had a greater discriminatory power than *ipTM* in predicting known true complexes from mismatched pairs of proteins in biosynthetic gene clusters (33).

### Usage and Output

The code is written in Python3 and takes as input a json file from AlphaFold2 or AlphaFold3 and corresponding PDB-format or mmCIF-format files for the coordinates respectively. The commands to use are:

~~~
python ipsae.py <path_to_json_file> <path_to_af2_pdb_file> <pae_cutoff> <dist_cutoff>
python ipsae.py <path_to_json_file> <path_to_af3_cif_file> <pae_cutoff> <dist_cutoff>
~~~

For example:

~~~
python ipsae.py RAF1_KSR1_scores_rank_001_alphafold2_multimer_v3_model_4_seed_003.json \ RAF1_KSR1_unrelaxed_rank_001_alphafold2_multimer_v3_model_4_seed_003.pdb 15 15
python ipsae.py fold_raf1_ksr1_mek1_full_data_0.json fold_raf1_ksr1_mek1_model_0.cif 15 15
~~~

The output from the second command is given in Figure 10.

**Figure 10.**
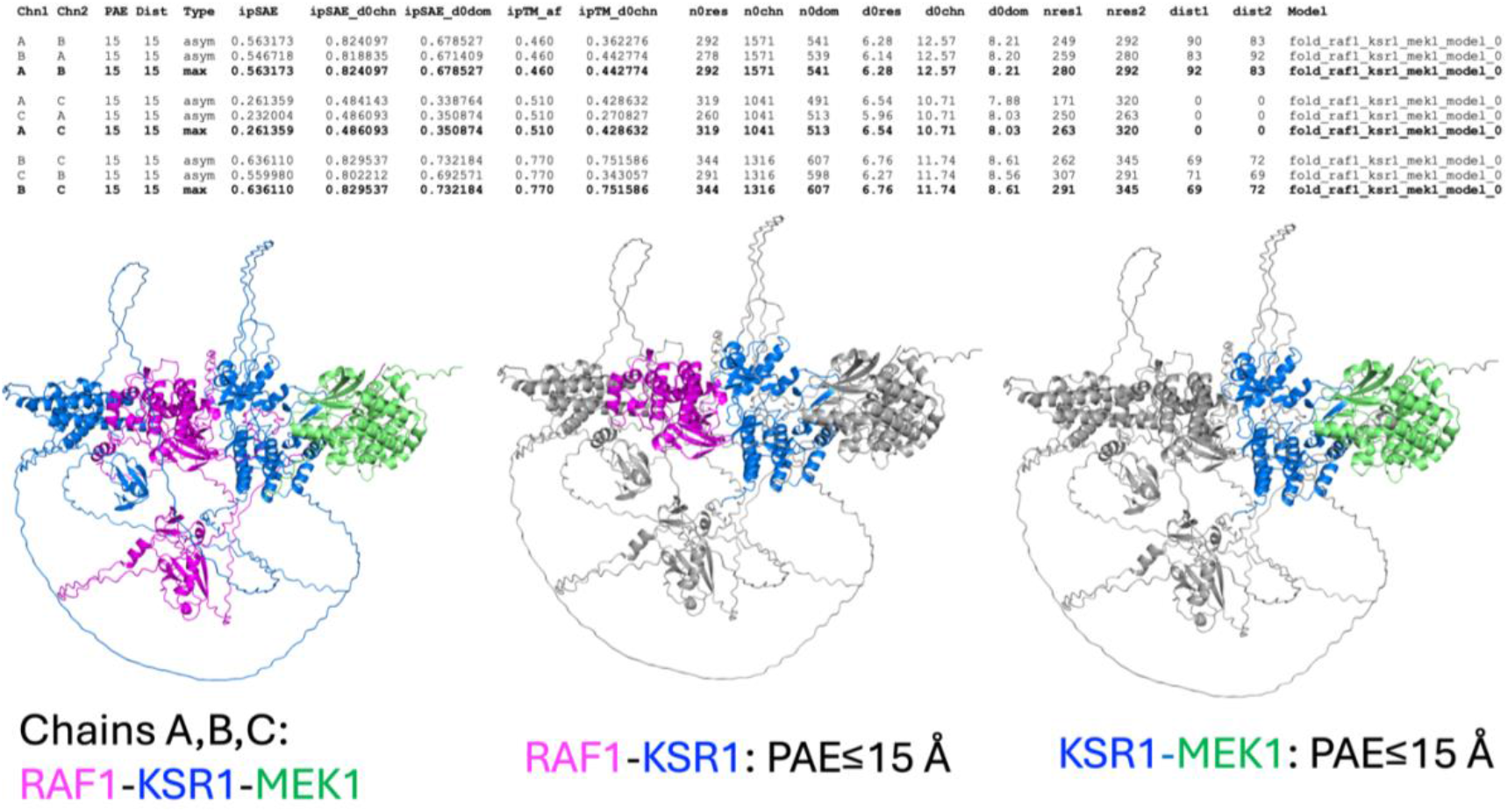
Output of ipsae.py on an AlphaFold3 model of a ternary complex of full-length human RAF1, KSR1, and MEK1 (Uniprots: RAF1_HUMAN, KSR1_HUMAN, MP2K1_HUMAN). The asymmetric values of the *ipSAE* metrics are given in rows with type equal to “asym.” The maximum value of each metric (over X⟶Y and Y⟶X asymmetric values) is given in the row labeled “max” (shown in bold type). Bottom: (left) top ranked AlphaFold3 model with chains labeled by color: RAF1 (chain A: magenta), KSR1 (chain B: blue), MEK1 (chain C: green). Middle: After coloring all three chains gray, PyMOL script alias “color_A_B” colors magenta and blue all residues in chains A and B respectively that have one or more interchain *PAE* values less than the cutoff (15 Å). Right: color B_C colors residues blue and green if they have interchain *PAE* values less than the same cutoff.

The code reads the overall *ipTM* from the AlphaFold2 json file, which has one value for any size protein complex. Given the name of the AlphaFold3 “full_data” json file, the code will read the *chain_pair_iptm* from the corresponding “summary_confidences” json file, if it exists. In this example, that would be named `fold_raf1_ksr1_mek1_summary_confidences_0.json` . In the output, these scores are called *ipTM_af*. For AlphaFold2, all chain pairs have the same value of *ipTM_af*. For the example in Figure 10, AlphaFold3 calculate *ipTM* pairwise values for 0.46 for RAF1-KSR1 (chains A and B), 0.51 for RAF1-MEK1 (chains A and C), and 0.77 for KSR1-MEK1 (chains B and C).

To calculate the *ipSAE* and other metrics, the code reads the *PAE* values from the respective json files. AlphaFold2 provides a square *PAE* matrix with row and column dimensions of the length of the combined protein sequences. The rows are aligned residues and the columns are scored residues. AlphaFold3 structure predictions may include post-translationally modified amino acids as well as ligands. The standard amino acids have single tokens and therefore single rows or columns in the *PAE* matrix in the json file. Modified amino acids, however, have one token per atom (e.g., phosphoserine, residue type SEP, has 10 tokens). We use the Cα atom as the appropriate token for the *PAE* matrix, so we can construct a square *PAE* matrix covering only one row or column per amino acid (whether modified or not). Ligands are excluded (label_seq_id=“.”).

Since AlphaFold2 does not calculate pairwise *ipTM* scores for multi-protein complexes and AlphaFold3 provides only the symmetric (maximum) pairwise *ipTM* scores, we use the *PAE* matrix to calculate pairwise *ipTM* scores. To calculate *d*_0_ for the *ipTM* calculation, we use the sum of the full-length protein sequences for each sequence pair, as AlphaFold2 does (for dimer complexes) and AlphaFold3 does for all chain pairs. This metric is called *ipTM_d0chn*, where *d0chn* indicates that *d*_0_ is calculated from the chain lengths. The values for the complex in Figure 10 are 0.443, 0.429, and 0.752 respectively (compare the *ipTM_af* values of 0.46, 0.51, 0.77 respectively).The small differences arise from using the *PAE* values in the pairwise *pTM* matrix (Equation 7), instead of the expectation value over the probability distribution of *PAE* (Equation 6).

With the *PAE* matrix and *PAEcutoff* value, we can calculate the asymmetric *ipSAE* scores (Equation 14) and the overall *ipSAE* score (Equation 16), which is the maximum value of the two asymmetric scores for each chain pair. For the regular *ipSAE* score, we use *d*_0_ based on the number of residues in the scored chain that have *PAE*<*PAEcutoff*, given the aligned residue in the aligned chain. The number of residues (*n0res*) and the value of *d*_0_ (*d0res*) are given in the output. For the example in Figure 10, the *ipSAE* values are: 0.563, 0.261, and 0.636. The columns *nres1* and *nres2* provide the number of residues in the first and second chains that have interchain *PAE* values (for the same pair of chains) less than the cutoff (15 Å in this case). In the asym lines, these are for the aligned residues and scored residues respectively. In the “max” lines, they are the maximum of the two asymmetric values. Thus, RAF1 and KSR1 have maximum values (scored or aligned) with *PAE* less than the cutoff of 280 and 292 residues respectively.

The next two columns, *dist1* and *dist2*, provide the number of residues with *PAE* less than the cutoff and Cα-Cα distance less than the distance cutoff set by the user (15 Å in this case). RAF1 does not contact MEK1 in the model, and the *dist1* and *dist2* values are both 0 for chains A+C. The *ipSAE* value is correspondingly only 0.261, while the *ipTM_af* value is 0.51 (probably because RAF1 can also interact with MEK1 but does not do so in this model).

The `ipsae.py` script outputs a PyMOL script with aliases to color residues in each pair of chains with *PAE* less than the cutoff. These residues are highlighted in magenta and blue in the middle structural figure in Figure 10 for RAF1+KSR1 and the right-side structural figure in blue and green for KSR1+MEK1.

The script also calculates two other forms of *ipSAE* for comparison purposes: *ipSAE_d0chn* and *ipSAE_d0dom. ipSAE_d0chn* uses the same *PAE* cutoff as *ipSAE* but calculates *d*_0_ from the sum of the two full-length sequence lengths (*n0chn, d0chn*). *ipSAE_d0dom* uses a value of *d*_0_ from the number of residues in the two chains that have any interchain *PAE* values less than the *PAEcutoff* (*nres1, nres2*).

Finally, for plotting figures like Figures 5 and 6 (e.g., the residue- and chain-pair specific values for *ipSAE*), the script outputs a file with name like:

~~~
fold_raf1_ksr1_mek1_model_0_15_15_byres.txt
~~~

with columns:

~~~
i, AlignChn, ScoredChain, AlignResNum, AlignResType, AlignRespLDDT, n0chn, n0dom, n0res, d0chn, d0dom, d0res, ipTM_pae, ipSAE_d0chn, ipSAE_d0dom, ipSAE.
~~~

The value *i* is the residue number across all chains (from 1 to total number of residues in model). The aligned chain refers to the chain with residues *i* in the pTM expressions, the scored chain covers residues *j. n0res* and *d0res* are residue-specific values for the number of residues with *PAE* less than the chosen cutoff and the corresponding *d*_0_ value. The other values are all chain-pair specific.

## Code availability

A python3 script is available at github.com/dunbracklab/IPSAE.

## Acknowledgments

I thank my lab members and members of the Fox Chase Cancer Center Molecular Modeling Facility for helpful discussions, including Mark Andrake, Sven Miller, Qifang Xu, Joan Gizzio, Pragya Priyadarshini, Brianna Trankle, and Xiyao Long. This work was funded by NIH grants R35 GM122517 (R.L.D.) and P30 CA006927 (Fox Chase Cancer Center). I thank Luca Genz and Maya Topf for running their benchmark on structure prediction accuracy of protein complexes and providing me with the results, as well as permission to use the figure they provided (Fig. 9).

